# Interpretable prognostic modeling of endometrial cancer

**DOI:** 10.1101/2022.08.23.504935

**Authors:** Bulat Zagidullin, Annukka Pasanen, Mikko Loukovaara, Ralf Bützow, Jing Tang

**Affiliations:** Research Program in Systems Oncology, Faculty of Medicine, University of Helsinki, 00290 Helsinki, Finland; Institute for Molecular Medicine Finland (FIMM), HiLIFE, University of Helsinki, 00290 Helsinki, Finland; Department of Pathology, University of Helsinki and Helsinki University Hospital, Helsinki 00290, Finland; Department of Obstetrics and Gynecology, Helsinki University Hospital and University of Helsinki, 00290 Helsinki, Finland; Research Program in Applied Tumor Genomics, Faculty of Medicine, University of Helsinki, 00290 Helsinki, Finland

## Abstract

Endometrial carcinoma (EC) is one of the most common gynecological cancers in the world. In this work we apply Cox proportional hazards (CPH) and optimal survival tree (OST) algorithms to the retrospective prognostic modeling of disease-specific survival in 842 EC patients. We demonstrate that the linear CPH models are preferred for the EC risk assessment based on clinical features alone, while the interpretable, non-linear OST models are favored when patient profiles are enriched with tumor molecular data. By studying the OST decision path structure, we show how explainable tree models recapitulate existing clinical knowledge prioritizing L1 cell-adhesion molecule and estrogen receptor status indicators as key risk factors in the p53 abnormal EC subgroup. We believe that visually interpretable tree algorithms are a promising method to explore feature interactions and generate novel research hypotheses. To aid further clinical adoption of advanced machine learning techniques, we stress the importance of quantifying model discrimination and calibration performance in the development of explainable clinical prediction models.

## Introduction

Endometrial carcinoma (EC) is the most common gynecologic malignancy in the OECD member states. In 2020, 417,000 new cases and 97,370 deaths have been attributed to the EC worldwide, which is a 10% increase in incidence and an 8% increase in mortality since 2018. Both metrics vary considerably geographically and across patients’ socioeconomic strata [1,2]. In the UK, the expected 5-year survival is 77%, with 85% for stage I disease and 25% for stage IV 3. EC treatment options depend on tumor staging and histological findings, which are prone to misdiagnosis [4]. Addition of molecular profiling information to histological features has been shown to improve patient stratification and subsequent selection of adjuvant therapies [5–9]. To further improve the EC risk assessment, it is important to develop transparent computational models that utilize both clinical and molecular patient profiles.

Two commonly used statistical methods in the survival analysis of EC patients are the Kaplan-Meier method and the Cox proportional hazards (CPH) regression. The Kaplan-Meier method is used to approximate cumulative survival probability (survival function) from lifetime and censored data [10]. It is well-suited to summarize survival functions from full cohorts, and it allows for their visual analysis. The CPH regression is the most popular model for the analysis of survival data when multiple variables are available [11]. Its utility is limited due to the CPH assumptions, such as the linearity and additivity of predictor variables, as well as the methodological difficulties related to variable selection. Machine learning (ML), such as deep learning and ensemble models, improve on these shortcomings. They have been shown to perform particularly well with high-dimensional datasets, such as -omics readouts, electronic health records, and high content imaging [12,13]. Deep learning and ensemble ML models have also been applied to prognostic prediction modeling of patient outcomes in the EC [14–17]. However, these ML models still see limited use in the clinical practice [18]. Their poor adoption may be attributed to the black-box nature that complicates model interpretability, a high risk of bias, and the need for larger training datasets to achieve similar performance, as compared to linear Cox regression [19].

Tree-based ML methods have been used to account for non-linear effects and variable interactions in survival analysis [20]. Tree-based ML methods are interpretable by design as every prediction made by a trained model can be associated with a corresponding decision path, and the hierarchical structure of the model as a whole can be easily visualized [21]. Further, they can take into account factors that may act differently in patient subgroups, unlike linear models that favor global factors with uniform effects across entire patient cohorts [22]. There are several variants of decision trees that can be used to estimate patient risks, such as the CART model proposed by Breiman et al or the conditional inference tree model by Hothorn et al [23,24]. While decision trees can be ensembled leading to better performance than single trees, like in the random survival forest algorithm by Ishwaran et al, this makes them considerably less interpretable [25–27]. In light of recent research advances aimed at improving decision tree algorithms through better splitting and pruning criteria, single decision tree models are a good alternative to the CPH regression in the development of explainable clinical prediction models [28,29].

In this retrospective study we explore a cohort of 842 EC patients with 43 clinicopathological and molecular features collected at the Helsinki University Hospital between 2007 and 2012. We report two interpretable models that predict disease-specific survival: a multivariable CPH regression and a visually interpretable optimal survival tree (OST) [29]. Both are built on two sets of variables: a clinical set and an extended set, which is enriched with molecular information of the EC patients, namely L1CAM (CD171) and estrogen receptor (ER) status indicators, as well as the cell cytology and tumor size. We use Harrell’s time-independent concordance index (C-index) and time-dependent integrated Brier score (IBS) to compare their performance [30,31]. These two measures report related, but distinct performance metrics, as C-index quantifies discrimination, or how well a model separates low-risk from high-risk patients, while IBS also quantifies calibration, which is the extent of an agreement between observed outcomes and model predictions [32]. In this work we show that to select an optimal EC prognostic model, a discrimination measure should be supplemented with a calibration measure, such as IBS [33–36]. We find that the CPH models trained on the clinical variables have better C-index than the OST models, whereas the IBS scores of both model types are comparable. Extending clinical data with tumor molecular profiles improves the discrimination and calibration performance in both model types, with a bigger improvement and overall best C-index and IBS values seen in the OST models. Finally, we suggest that the Cox proportional hazards regression should be used in the EC risk assessment based on clinical data only, while optimal survival trees are preferred when molecular information is available.

## Materials and Methods

### Study cohort

This retrospective analysis is based on a cohort of 842 patients with unselected EC that underwent surgical treatment between 2007 and 2012 at the Helsinki University Hospital. The follow-up time ranges from 1 to 136 months with a median of 82 months. In total, 591 (70.2%) patients survived until the end of the study, 148 (17.6%) died from the EC, 103 (12.2%) died from other causes. The endpoint of interest is disease-specific survival (DSS). Based on tumor molecular profiles derived through The Cancer Genome Atlas project, 604 (71.7%) patients were assigned to one of four ProMisE classes, for the remaining 238 (28.2%) patients the ProMisE categories were not assigned experimentally [5,6]. Four categories are: a) mismatch repair deficient (MMRd), b) no specific molecular profile (NSMP), c) p53 abnormal and d) polymerase-ε hypermutated (POLE). Among 604 patients that have ProMisE classes assigned to them, 74 died due to other causes and 30 belong to the POLE subgroup, where no one died from the EC. Each patient is described with a feature vector consisting of 43 variables, out of which 33 are categorical and 10 are numeric. Please refer to the Supplementary Materials - Extended variable information for a more detailed variable description.

### Data preprocessing

All numeric variables, except for age and BMI, are winsorized at the 99% level to limit the effect of extreme values using the quantile function derived via the inverse of an empirical distribution function [37,38]. Variables with more than five categories, such as stage, or those with unbalanced class proportions, such as adjuvant therapy, are simplified by combining subcategories together.

We impute missing values to prevent the exclusion of observed data [39]. Missing values are imputed using the multivariate imputation by chained equations method, where numerical and binary variables are predicted with random forest models consisting of 100 decision trees, unordered categorical data with more than two levels are imputed with the polytomous regression, and ordered categorical variables with more than two levels are imputed with the proportional odds model [40]. Variables are imputed in the order of low to high proportion of missingness. R mice package version 3.14.7 is used to generate 120 imputed datasets, which are subsequently merged by taking mean values for the numeric variables and mode values for categorical variables [41]. The response variable is kept throughout the imputation [42].

Finally, to select variables for the CPH regression models we compare the distributions of numerical and binary categorical variables, stratified by the response. We apply the Pearson correlation coefficient to identify collinear numerical variables, and Goodman and Kruskal’s lambda to identify associated categorical variables [43]. Our primary goal is to optimize data for the CPH regression performance. Therefore, simplification of categorical data and variable selection in the subsequent steps are iterated several times. We use the analysis of deviance test to compare nested CPH models, while the Akaike information criterion is preferred for the comparison of non-nested models [44,45].

The complete experimental pipeline is shown in Figure 1.

**Figure 1.**
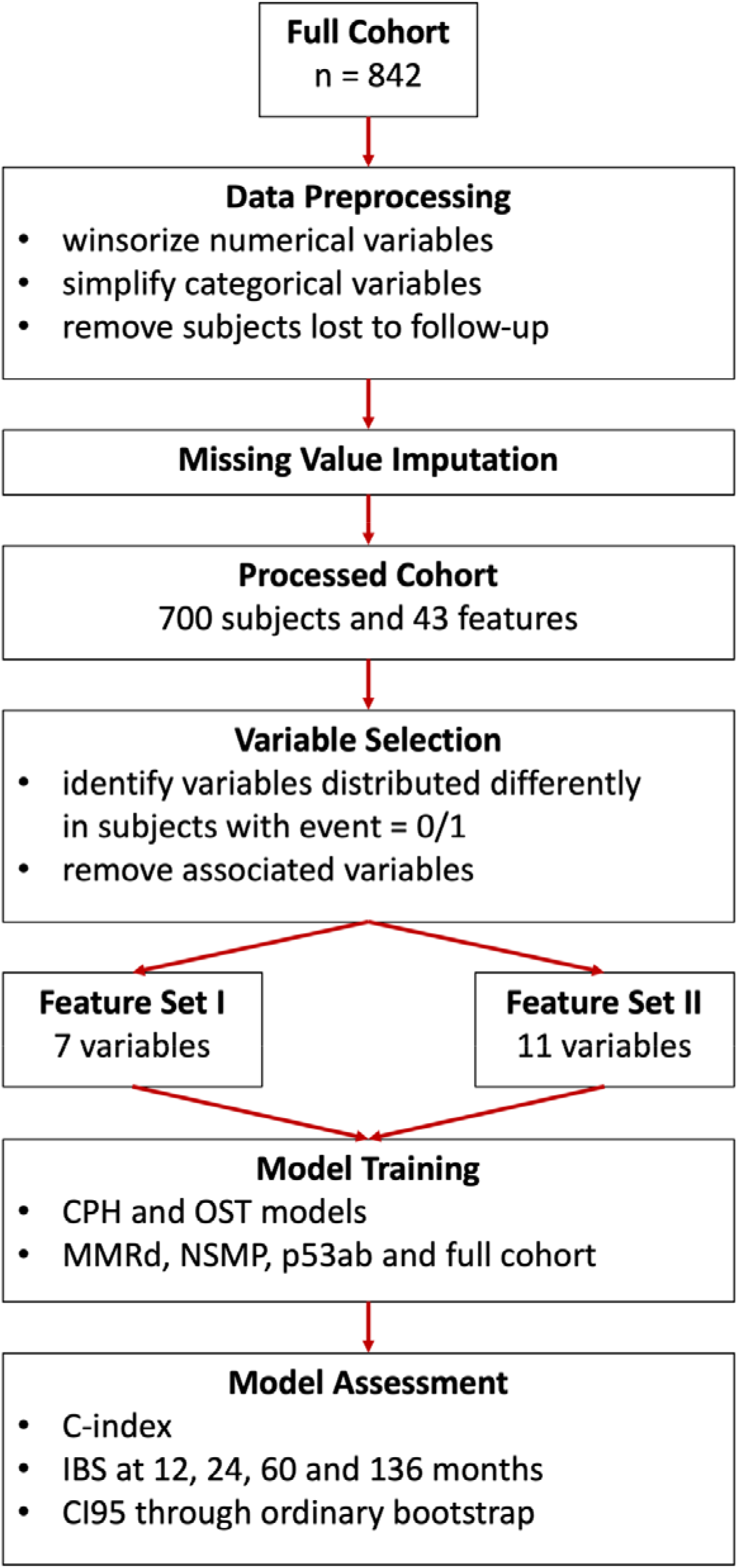
Experimental pipeline.

### Statistical modeling

We train two types of interpretable models to predict individual survival probabilities for the full patient cohorts, and subcohorts stratified by the ProMisE classes. We assess their performance using C-index and IBS. We estimate 95% confidence intervals for the performance metrics by 1000 repetitions of the ordinary bootstrap with replacement [46]. We also report the performance of seven additional survival analysis models using the C-index metric in the Supplementary Materials - Additional ML models section. We follow the Transparent Reporting of a multivariable prediction model for Individual Prognosis Or Diagnosis (TRIPOD) Initiative to enable interoperability of the developed models and to decrease reporting bias [47].

### Cox proportional hazards model

Survival CPH regression is defined as a product of a non-parametric hazard function *λ*(*t*) and the *e^ß^* term, where *t* is time, *X* is a vector of variables describing a patient, and *β* is a vector of the model’s coefficients. The *λ*(*t*) part of the CPH model is identical for all patients at a time *t*. It is referred to as a hazard function of a standard patient, which is a patient with *Xβ*=0. The second term is patient-specific, and it is used to calculate a hazard ratio without knowing the hazard function *λ*(*t*), where the hazard ratio is the risk of death in relation to a control [48]. We use the Breslow method, as implemented in R riskRegression package version 2022.03.22, to specify the hazard function, which is required for estimating individual survival probabilities [49]. We use Schoenfeld residuals to test for the proportional hazards assumption, as implemented in R survival package version 3.3.1. We estimate CPH model parameters by maximizing the partial log-likelihood.

### Optimal survival tree model

We use the optimal survival tree (OST) method to develop interpretable decision tree models for estimating patient survival probabilities. The OST algorithm creates multiple candidate decision trees and optimizes their variable splitting thresholds one variable at a time using coordinate descent [50]. The main idea is to use previously optimized parameters in subsequent splitting criteria updates, ultimately outputting a single decision tree that can be visually examined. The OST loss function compares how close the predicted *e^ß^* terms for each patient are to the cumulative survival probabilities, obtained by the Nelson-Aalen estimator [29]. We prioritize model robustness in the training process by: a) limiting the tree size, since too deep or too wide trees obfuscate the model interpretability, b) increasing the number of random restarts to use in the local search algorithm, and c) controlling the minimum number of points that must be present in every leaf node of the fitted trees. The complexity parameter that determines the tradeoff between the accuracy and model complexity is tuned automatically by assessing the out-of-sample performance. The patient cohorts for the OST model training are split, such that the complete cases are used for model fitting, and the imputed subsets are used for validation. The final validated models are then retrained on the combined (complete case and imputed) patient cohorts. We fit the OST models, as implemented in R iai package version 1.7.0, using the log-likelihood criterion [51].

### Model performance metrics

C-index reports model discrimination performance, i.e. the model’s ability to predict correct rankings of the survival times. C-index is defined as a ratio of concordant pairs of subjects to the total number of comparable pairs. A pair is concordant when a subject with shorter survival time is estimated to have a higher risk than the one with longer survival time. A pair is comparable if a) it is possible to determine which subject experienced the event first or b) a subject with a shorter survival time experienced an event, while the other one is censored and is not lost to follow-up yet. C-index ranges between 0 and 1, where higher values are better.

IBS reports both model discrimination and calibration, i.e. the extent of an agreement between observed outcomes and model predictions [52,53]. Brier score is defined as a mean squared difference between event indicators and predicted survival probabilities at a time *t* [31]. By summing Brier scores over a time interval we obtain the integrated Brier score (IBS), which is then adjusted for patients lost to follow-up using the inverse probability censoring weighting method [54]. We use R pec package version 2022.05.04 to compute IBS at 12, 24, 60, and 136 months based on the predicted individual survival probabilities of patients. IBS ranges between 0 and 1, where lower values are better.

### Computational resources

All computations are performed using R 4.2.0 on MacOS 12.5 and Python 3.9.7 on Ubuntu 20.04 LTS.

## Results

The initial cohort consists of 842 patients diagnosed with unselected endometrial carcinoma. Following missing value imputation, excluding subjects that died due to other causes and those that belong to the POLE group, where no one died, the final analysis cohort consists of 700 patients. Among those 700 patients, 305 (43.6%) belong to the MMRd subgroup, 308 (44%) belong to the NSMP subgroup and 87 (12.4%) belong to the p53ab subgroup. Majority of the tumors are histopathological grade 1-2 (74%) and stage I disease (73%). The median follow-up time of censored cases is 92 (interquartile range, 78-122) months. There are 182 subjects who had disease recurrence and 147 died during the follow-up time. Patient demographics are shown in Table 1.

**Table 1.**
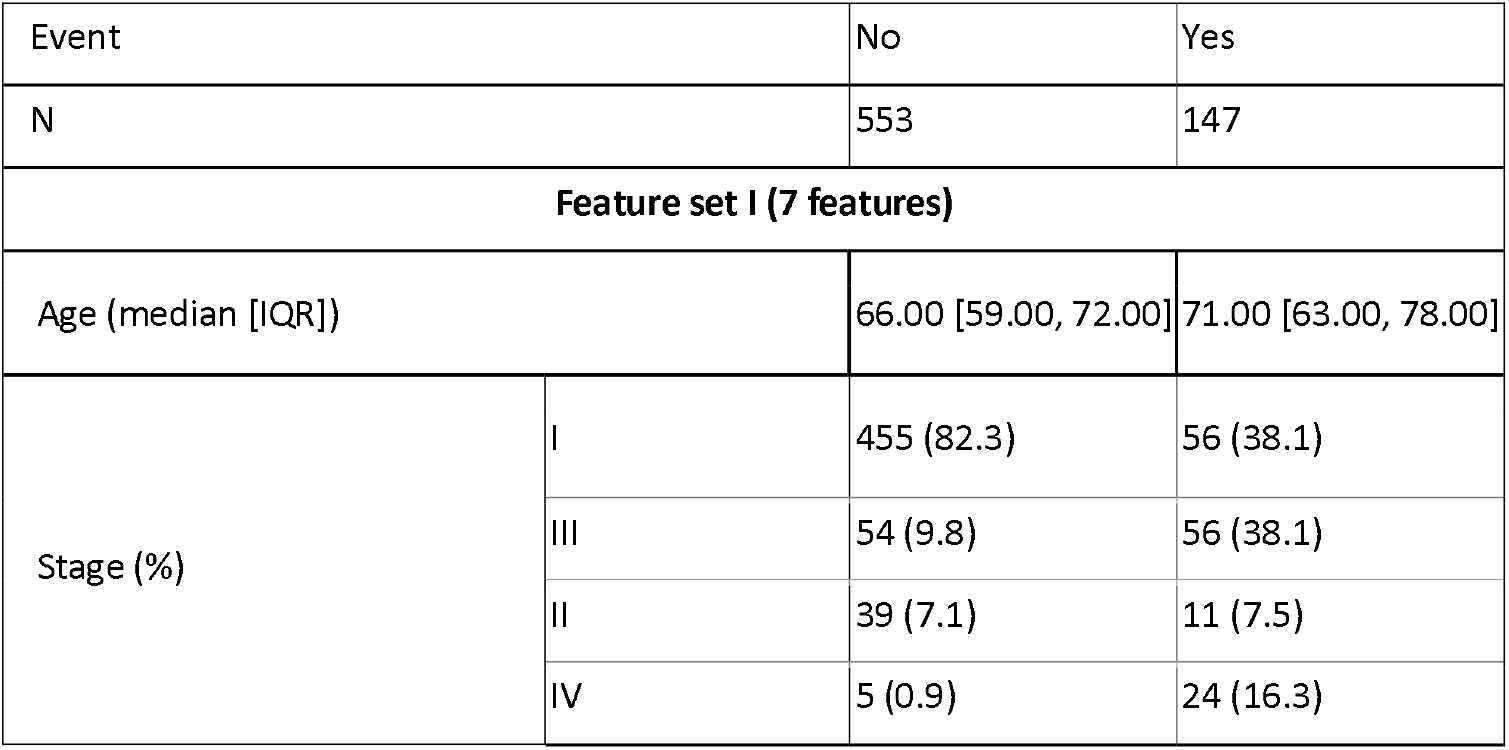
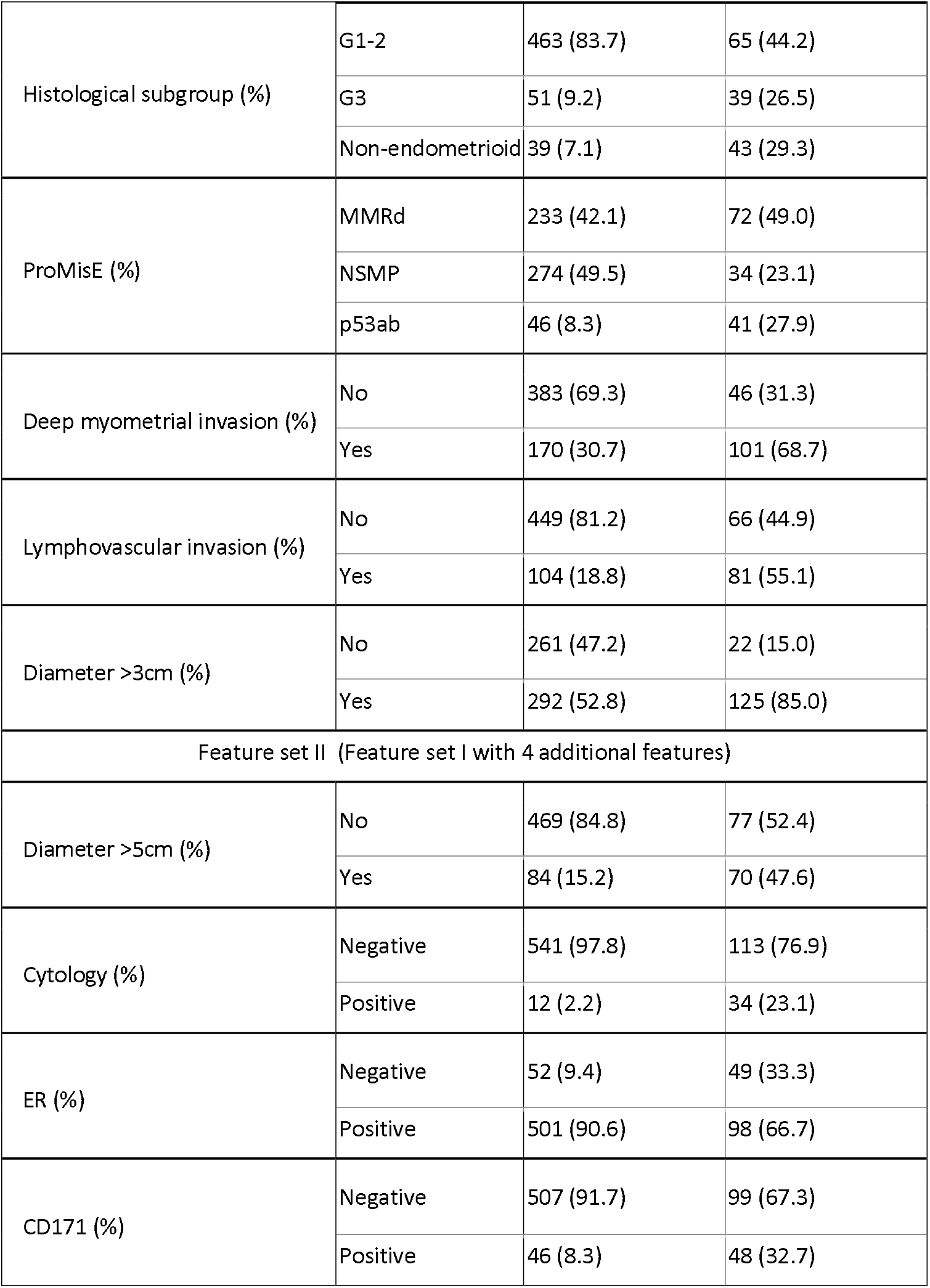
Patient demographics (n = 700). Feature set I consists of 7 features (FSI), and feature set II consists of 11 features (FSII).

The multivariable CPH models are compared with the OST models in prediction of the DSS using two feature sets in four patient cohorts. Variable selection for both feature sets is performed to optimize the CPH model performance. Subsequently, the OST models are fit on the selected feature sets. The feature set I (FSI) consists of seven variables: age, stage, histological subgroup, ProMisE, deep myometrial invasion, lymphovascular invasion, and tumor diameter > 3cm. The feature set II (FSII) adds four more variables to the FSI, namely tumor diameter > 5cm, cytology, ER status, and postoperative L1CAM (CD171) expression.

### Model discrimination

The C-index scores of the CPH and OST models with the corresponding 95% confidence intervals are shown in Table 2.

**Table 2.**
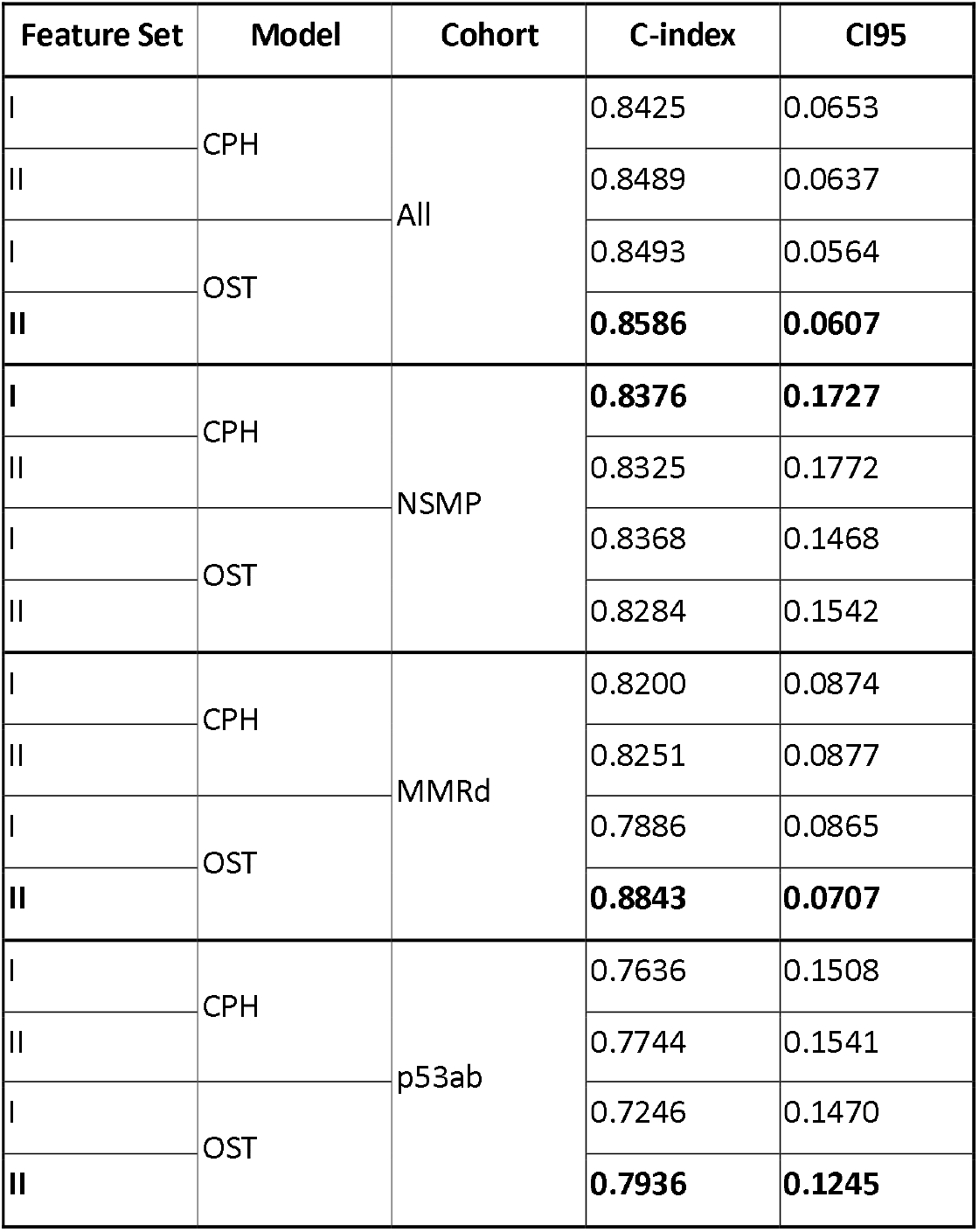
C-index of the Cox proportional hazards (CPH) models vs optimal survival tree (OST) models using. Two feature sets are: FSI (7 features) and FSII (11 features). Models in bold perform the best in their corresponding cohorts. 95% confidence intervals (CI95) are calculated using 1000 iterations of the ordinary bootstrap with replacement.

| Feature Set | Model | Cohort | C-index | CI95 |
| --- | --- | --- | --- | --- |
| I | CPH | All | 0.8425 | 0.0653 |
| II |  |  | 0.8489 | 0.0637 |
| I | OST |  | 0.8493 | 0.0564 |
| II |  |  | **0.8586** | **0.0607** |
| I | CPH | NSMP | **0.8376** | **0.1727** |
| II |  |  | 0.8325 | 0.1772 |
| I | OST |  | 0.8368 | 0.1468 |
| II |  |  | 0.8284 | 0.1542 |
| I | CPH | MMRd | 0.8200 | 0.0874 |
| II |  |  | 0.8251 | 0.0877 |
| I | OST |  | 0.7886 | 0.0865 |
| II |  |  | **0.8843** | **0.0707** |
| I | CPH | p53ab | 0.7636 | 0.1508 |
| II |  |  | 0.7744 | 0.1541 |
| I | OST |  | 0.7246 | 0.1470 |
| II |  |  | **0.7936** | **0.1245** |

Model discrimination performance is improved by the inclusion of additional variables, as indicated by higher C-index scores in the FSII versus FSI feature sets. The OST models trained on the FSII report the highest overall C-index in all the subcohorts, except for the NSMP ProMisE class, where the CPH model trained on FSI has the highest C-index of 0.8376, followed by the OST model with the C-index of 0.8368. We note that the CPH models report on average 2.2% higher C-index than the OST models in the FSI. This trend is reversed in the FSII, where the OST models report on average 2.5% better C-index than the CPH models. The largest C-index increase for the former is 10.8% in the MMRd and 8.7% in the p53ab subcohorts, while the largest increase for the latter is 1.4% in the p53ab subcohort. Overall, non-linear optimal survival tree models benefit more from the addition of molecular information than the linear Cox proportional hazards models.

### Model calibration

We report the IBS scores with the 95% confidence intervals for both the CPH and the OST models at 12, 24, and 60 months, and the overall IBS at 136 months of follow-up in Figure 2 and Supplementary Table 1.

**Figure 2.**
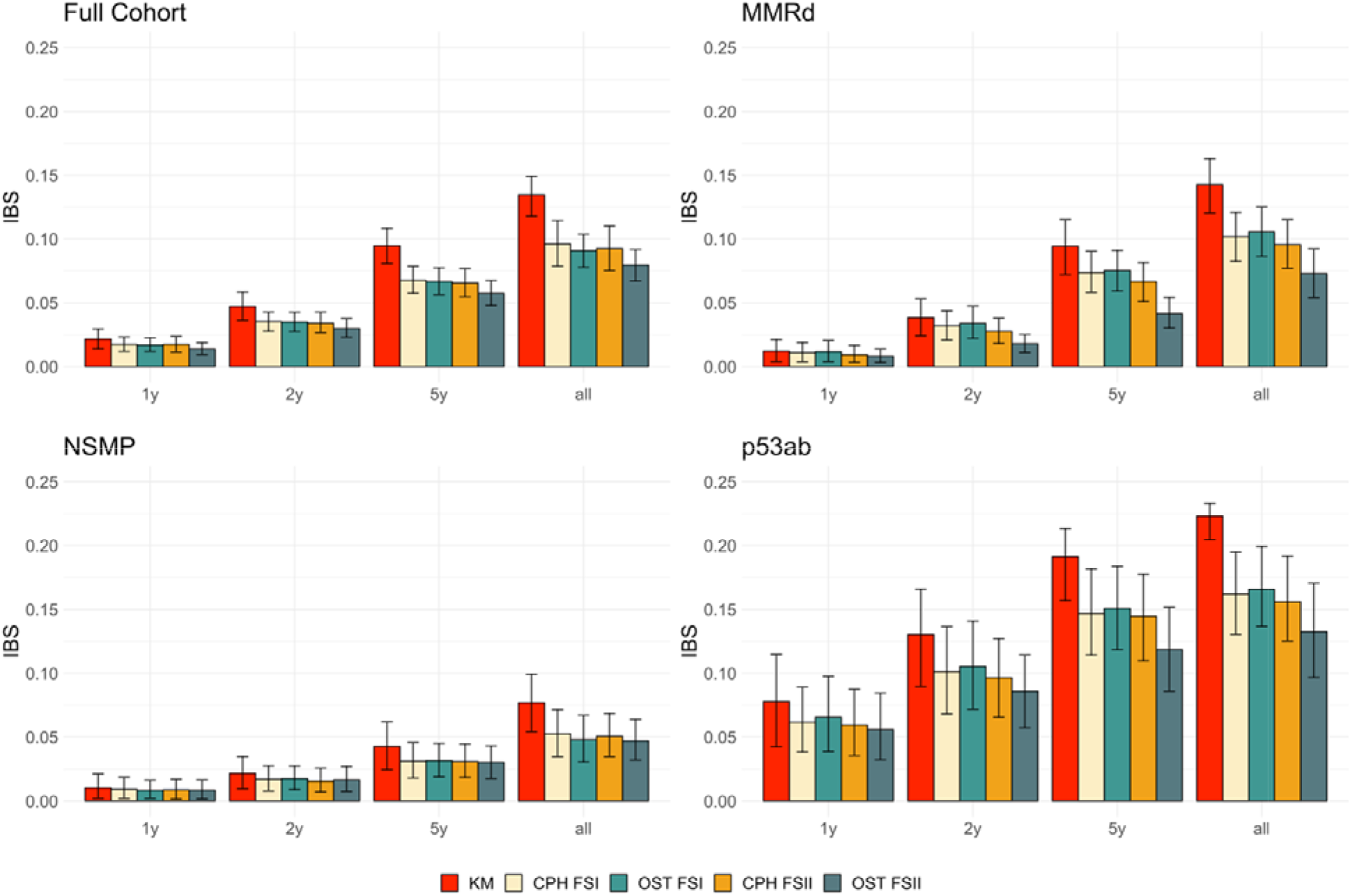
Integrated Brier score (IBS) at 1 year, 2 years, 5 years and 136 months (all) for models trained on four patient cohorts. KM is a non-parametric Kaplan-Meier estimator that may be used as a reference for the parametric models. Cox proportional hazards (CPH) and optimal survival tree (OST) models are trained on two feature sets: FSI with 7 features and FSII with 11 features. Error bars indicate 95% confidence intervals calculated using ordinary bootstrap with replacement, repeated 1000 times.

All models across all cohorts and feature sets show better (i.e. lower) IBS at shorter follow-up times, e.g. the IBS scores at 1 year of follow-up are up to an order of magnitude lower than at 136 months of follow-up. Both OST and CPH model types generally report better IBS scores when trained on a feature set with molecular information (FSII), as compared with the models trained on clinical measurements (FSI). The OST models improve more from the additional molecular features than the CPH models. The OST models trained on the FSII report 15.4% better IBS at 1 year, 21.6% better IBS at 2 years, 21.0% better IBS at 5 years and 16.2% better IBS at the complete follow-up, as compared with the FSI-trained OST models. The IBS improvements for the CPH models trained on the FSII are 5.7% at 1 year, 7.5% at 2 years, 3.9% at 5 years and 4.4% at the complete follow-up, as compared with the FSI-trained models. Both model types are on par with each other on the FSI feature set, however, the OST models have better IBS scores than the CPH models on the FSII set.

### Model interpretation

The hazard ratios with the corresponding 95% confidence intervals of the CPH models trained on a full cohort on two feature sets are in Table 3. It is important to note that the interpretation of the CPH model coefficients should be performed when the proportional hazards (PH) assumption is satisfied. We found evidence that the terms “non-endometrioid” of the histological subgroup and ER+ status do not satisfy the PH assumption in the CPH models built on the FSI or FSII feature sets according to the Schoenfeld residual test. Upon the visual inspection, the violations are minor for both. Further, since both model types pass the global PH test with the p-values of 0.245 and 0.25, respectively, we deem it appropriate to ignore the PH violations.

**Table 3.** Hazard ratios (HR) of the Cox proportional hazards model trained on the full cohort using 7 features (FSI) and 11 features (FSII) with Wald 95% confidence intervals and log-rank test p-values.

| Term | HR on FSI | p-value | HR on FSII | p-value |
| --- | --- | --- | --- | --- |
| Age | 1.04 (1.02-1.06) | 1.78E-05 | 1.04 (1.02-1.06) | 9.73E-06 |
| Stage II | 1.33 (0.68-2.58) | 4.08E-01 | 1.33 (0.68-2.61) | 4.07E-01 |
| Stage III | 2.73 (1.78-4.19) | 4.06E-06 | 2.2 (1.4-3.47) | 6.80E-04 |
| Stage IV | 7.85 (4.35-14.16) | 7.52E-12 | 3.81 (1.9-7.63) | 1.64E-04 |
| ProMisE MMRd | 1.61 (1.05-2.47) | 3.07E-02 | 1.8 (1.17-2.77) | 7.77E-03 |
| ProMisE p53ab | 2 (1.2-3.32) | 7.57E-03 | 1.88 (1.13-3.14) | 1.51E-02 |
| Histological subgroup G3 | 2.04 (1.32-3.16) | 1.29E-03 | 1.66 (1.05-2.63) | 2.99E-02 |
| Histological subgroup Non-endometrioid | 1.35 (0.82 - 2.21) | 2.40E-01 | 0.95 (0.54-1.68) | 8.70E-01 |
| Deep myometrial invasion Yes | 1.25 (0.83 - 1.88) | 2.89E-01 | 1.13 (0.74-1.73) | 5.69E-01 |
| Lymphovascular invasion Yes | 1.94 (1.35 - 2.79) | 3.45E-04 | 2.05 (1.42-2.97) | 1.25E-04 |
| Diameter >3cm Yes | 2.35 (1.44 - 3.83) | 6.49E-04 | 2.35 (1.37-3.88) | 1.39E-03 |
| Diameter >5cm Yes |  |  | 1.29 (0.88-1.9) | 1.89E-01 |
| Cytology Positive |  |  | 2.73 (1.68-4.43) | 4.95E-05 |
| ER Positive |  |  | 0.7 (0.45-1.09) | 1.14E-01 |
| CD171 Positive |  |  | 1.37 (0.97-2.17) | 1.78E-01 |

Age, more advanced disease stages, larger tumor sizes, deep myometrial invasion, lymphovascular space invasion, positive cytology, ER- and L1CAM+ terms are all associated with worse survival outcomes [55,56]. The MMRd and p53 aberrant EC classes are identified as more aggressive EC forms than the NSMP class, with the HR of 1.61 and 2 (1.8 and 1.88 in the FSII), respectively. Similarly, histological subgroup G3 leads to a higher risk of death than the G1-G2 subgroup with the HR of 2.04 on the FSI and 1.66 on the FSII. Interestingly, the non-endometrioid EC subtype is not robustly associated with a higher risk according to the HR values of 1.23 and 0.95 in FSI and FSII feature sets, respectively. This ambiguity in assessing the survival differences between type I and type II tumors has been previously reported in the literature [57,58].

We next explore how the tree models may supplement conventional linear methods in the interpretation of EC risk factors by studying the OST and CPH model types trained on the p53ab subcohort and the FSII feature set. We focus on the p53ab subgroup (n = 87), as it shows the largest relative improvement in the C-index from the addition of molecular information in the CPH models (0.7636 vs 0.7744) and the second largest in the OST models (0.7246 vs 0.7936). The CPH IBS values improve by 3% between FSII and FSI, whereas for the OST model the improvement is 19%. The HR scores with the 95% confidence intervals of the FSII-trained CPH model are in Table 4. The decision tree for the FSII-trained OST model is in Figure 3.

**Table 4.**
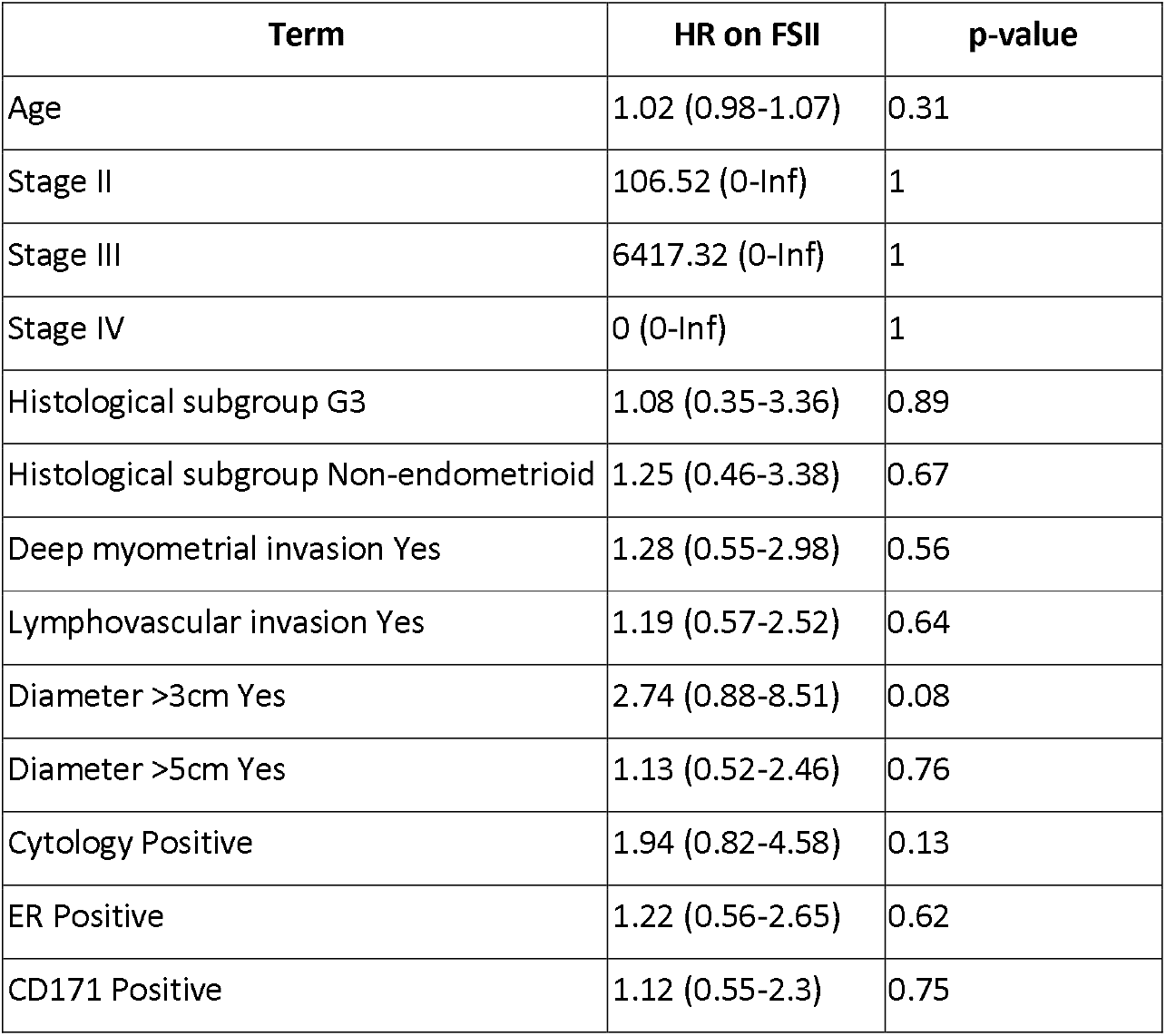
Hazard ratios (HR) of the p53ab subcohort Cox proportional hazards model trained on the FSII with Wald 95% confidence intervals and log-rank test p-values.

| Term | HR on FSII | p-value |
| --- | --- | --- |
| Age | 1.02 (0.98-1.07) | 0.31 |
| Stage II | 106.52 (0-Inf) | 1 |
| Stage III | 6417.32 (0-Inf) | 1 |
| Stage IV | 0 (0-Inf) | 1 |
| Histological subgroup G3 | 1.08 (0.35-3.36) | 0.89 |
| Histological subgroup Non-endometrioid | 1.25 (0.46-3.38) | 0.67 |
| Deep myometrial invasion Yes | 1.28 (0.55-2.98) | 0.56 |
| Lymphovascular invasion Yes | 1.19 (0.57-2.52) | 0.64 |
| Diameter >3cm Yes | 2.74 (0.88-8.51) | 0.08 |
| Diameter >5cm Yes | 1.13 (0.52-2.46) | 0.76 |
| Cytology Positive | 1.94 (0.82-4.58) | 0.13 |
| ER Positive | 1.22 (0.56-2.65) | 0.62 |
| CD171 Positive | 1.12 (0.55-2.3) | 0.75 |

**Figure 3.**
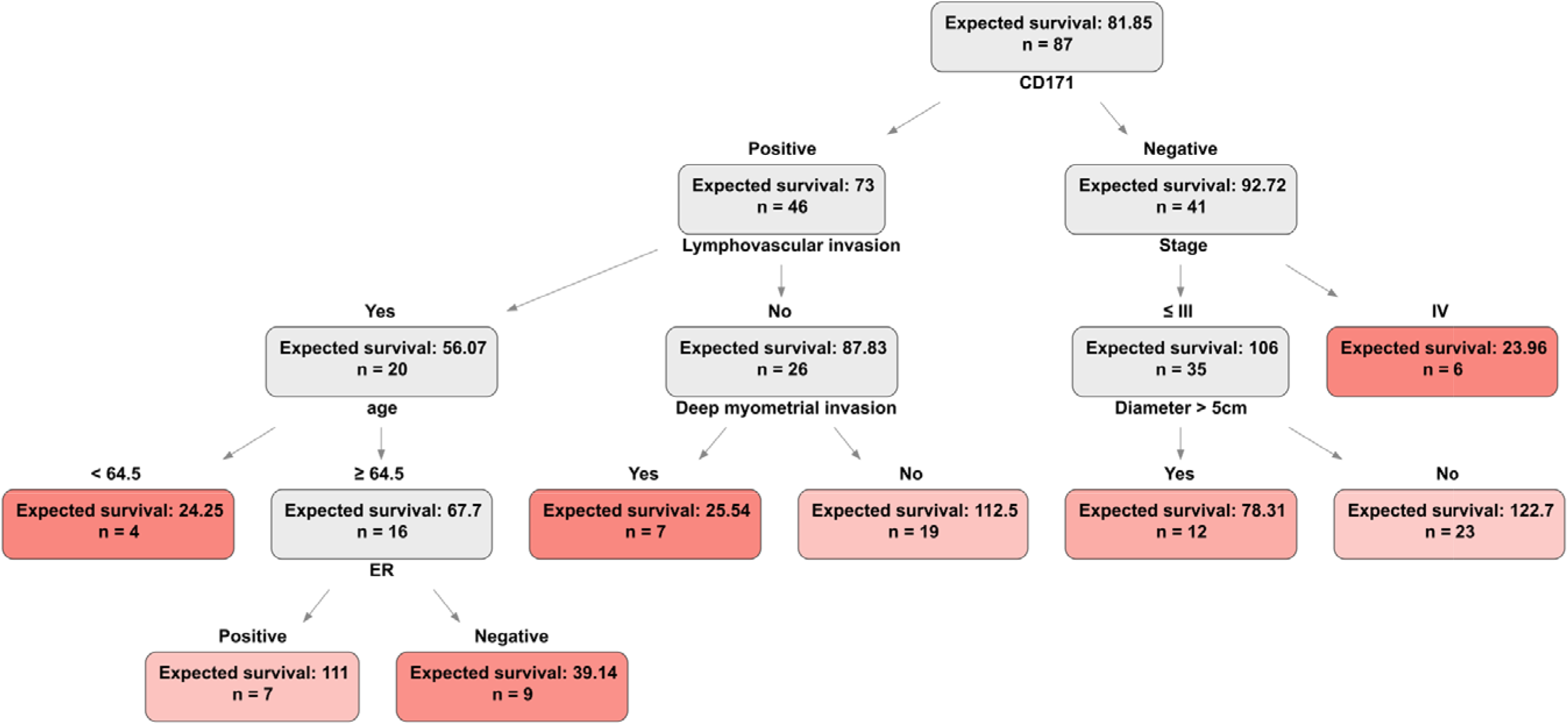
Optimal survival tree for the p53ab subcohort (n = 87) trained on the FSII set consisting of 11 features. Colors indicate leaf (terminal) nodes. Darker hues denote shorter expected survival measured in months and calculated via the integral of a survival function.

The CPH model reports a C-index of 0.7744, but some of the model coefficients are not informative nor do they support existing clinical evidence. For instance, the 95% confidence intervals of the HR values for the stage variable do not have an upper bound, positive ER status is indicated as a risk factor, and no p-values for the model coefficients are significant (Table 4). Therefore, it is not advised to use this model as is for the downstream tasks that require model interpretation. The OST model reports a C-index of 0.7936, and the decision tree path recapitulates some of the existing clinical knowledge (Figure 3). The L1CAM status is selected as the root node, i.e. the most informative variable to stratify the cohort on. The OST model correctly classifies the ER status as an important risk factor in the p53ab group. The clinical significance of ER and L1CAM in the non-endometrioid p53 aberrant tumors that are overrepresented in the p53ab ProMisE class (49.5% of subjects belong to the non-endometrioid subtype vs 11.7% in the full cohort) is supported in the literature [56,59,60]. This analysis demonstrates how tree-based ML is preferred to conventional Cox regression in the EC risk assessment when tumor molecular information is available.

## Discussion

We have trained the CPH and OST models on the full, MMRd, NSMP and p53ab patient cohorts using clinical and extended feature sets. Linear CPH and non-linear OST models trained on seven clinical variables report comparable discrimination performance with the C-index of 0.8425 vs 0.8493 in the complete cohort, and 0.8376 vs 0.8368 in the NSMP ProMisE class, respectively. Model calibration scores are also similar with a 5-year IBS of 0.0677 vs 0.0666 in the full cohort and 0.0309 vs 0.0314 in the NSMP class, respectively. In contrast, the CPH models have a better discrimination performance than the OST models with the C-index of 0.82 vs 0.7886 in the MMRd class, and 0.7636 vs 0.7246 in the p53ab class. The CPH models are as well-calibrated as the OST models in these subcohorts with the 5-year IBS of 0.0736 vs 0.0752 and 0.1467 vs 0.1508, respectively. According to the discrimination and calibration performance, the CPH models are preferred to the OST models in prognostic EC modelling using patient clinical data.

By enriching the clinical variables with patient molecular information, in particular the ER and L1CAM status indicators, and two histopathological features, namely cell cytology and tumor size <5cm we improve the discrimination and calibration performance of the CPH and OST models. The OST models make better use of additional features and are overall the best EC risk assessment models in the complete (C-index of 0.8586, IBS at 5 years of 0.0573), p53ab (C-index of 0.7936, IBS at 5 years of 0.1185) and MMRd cohorts (C-index of 0.8843, IBS at 5 years of 0.0416). Further, we show how interpretable OST decision trees may offer insights into the molecular mechanisms of the EC where the conventional CPH analysis falls short. The p53ab OST model trained on the extended feature set indicates that the L1CAM and ER status are important predictors in the non-endometrioid p53 aberrant tumors, while the p53ab CPH model assigns ER positive status (p-value of 0.62) as a risk factor contrary to the existing evidence, nor does it provide meaningful HR values for the disease stages. We conclude that when patient clinical data can be enriched with molecular measurements the OST method is preferred to the CPH regression in the EC risk assessment due to good discrimination and calibration performance, as well as the model interpretability through the decision path analysis.

There are several limitations in our study. Firstly, better prognostic survival models could be created if we had access to an external validation cohort [61,62]. In general, we hope that the research community could share anonymized patient datasets more freely, as openaccess initiatives contribute to the development of better prognostic prediction models [63]. Further, in addition to IBS, we are interested in exploring other model calibration measures, such as the integrated calibration index or standardized mortality ratio [64,65]. The third limitation stems from the methodological difficulties in the assessment of data imputation methods and their downstream effects. In this work we did not perform any formal tests to identify the missingness type, assuming missing at random for all explanatory covariates [66,67]. We performed an ad hoc assessment of imputation quality by comparing imputed variable distributions with those in the complete case cohorts. More robust and comprehensive methods for the assessment of data imputation techniques are needed [68].

## Conclusion

We show that the Cox proportional hazards and optimal survival tree models are well-suited for the prognostic survival modeling of endometrial carcinoma. The Cox proportional hazards regression is the method of choice for the EC risk assessment on the clinical feature set. Extending clinical variables with molecular tumor information, in particular the ER and L1CAM status indicators, improves the discrimination and calibration performance in both model types. Due to the overall best C-index and IBS scores and the interpretable structure, we recommend optimal survival tree models when tumor molecular data are available. Finally, we stress the importance of reporting model discrimination and calibration metrics to promote further adoption of ML prognostic models into the clinical practice.

## Additional Information

## Acknowledgments

We acknowledge the computational resources provided by the Finnish IT Center for Science. We acknowledge the University of Helsinki for the open-access fees.

## Author Contributions

BZ analyzed the data and wrote the manuscript. BZ, AP, MK and JT edited the manuscript. ML collected the clinical data. AP and RB reviewed patient histology, constructed the TMA used for immunohistochemical stainings and scored all the stainings. JT supervised the study. All authors have read and agreed to the published version of the manuscript.

## Institutional Review Board Statement

The study was conducted according to the guidelines of the Declaration of Helsinki.

## Informed Consent Statement

This study was approved by the Institutional Review Board of the Helsinki University Hospital (journal number 135/13/03/03/2013, date 29 May 2013) and the National Supervisory Authority for Welfare and Health (journal number 753/06.01.03.01/2016, date 9 February 2016). Patient consent was waived because of the retrospective nature of the research.

## Data Availability

The code and individual survival probabilities estimated using the OST and CPH models are available on https://github.com/netphar/survival_analysis. The raw patient data are not publicly available due to privacy restrictions but are available on reasonable request and with permission of clinical collaborators.

## Supplementary Materials

### Extended variable information

Each patient is described with a feature vector consisting of 43 variables, out of which 33 are categorical and 10 are numeric. Numeric variables are:

- Age;
- β-subunit of human chorionic gonadotropin (βhCG);
- Body mass index (BMI);
- Cancer antigen 125 (CA-125);
- Creatinine;
- Hemoglobin;
- Hematocrit;
- Human chorionic gonadotropin (hCG);
- Leukocytes;
- Thrombocytes.

Categorical variables are binary, unless stated otherwise:

- Adjuvant therapy (chemotherapy, vaginal brachytherapy, whole pelvic radiotherapy, whole pelvic radiotherapy with chemotherapy);
- ARID1a protein loss;
- CTNNB1 nuclear expression (negative, diffuse, focal);
- HER-2/neu expression (negative, diffuse and strong, diffuse and weak, focal);
- Histological subgroup (grade 1-2, grade 3, non-endometrioid);
- HNF1b positivity;
- Presence of endometrial hyperplasia;
- E-cadherin expression (negative, normal, weak);
- Estrogen (ER) and progesterone receptor (PR) expression;
- FIGO stage (I-II-III-IV);
- Final histology (carcinosarcoma, clear cell carcinoma, endometrioid carcinoma, serous carcinoma, undifferentiated carcinoma);
- Iliacal lymphadenectomy;
- KRAS mutation;
- Lymphadenectomy;
- p16 expression (negative, diffuse, focal);
- Paraaortic lymp node status;
- Peritoneal cytology;
- Postoperative Mayo criterion;
- Preoperative histology (low and high risk);
- Pre- and postoperative L1CAM (CD171) expression;
- ProMisE class (MMRd, NSMP, p53ab, POLE);
- Smoking status;
- Tumor infiltrating leukocytes (none, moderate, abundant);
- Tumor infiltrating leukocytes PD-L1 expression binarized at 1% and 10%;
- Uterine risk factors:

- Tumor diameter at 2cm/3cm/5cm levels;
- Deep myometrial invasion (≥50%);
- Lymphovascular space invasion;
- Myometrial invasion with levels <33%, 33%-66%, >66%;
- Vimentin expression (negative, diffuse, focal);

More detailed information regarding the study cohort is available in prior work 69,70.

### IBS model scores

**Supplementary Table 1.** IBS of Cox proportional hazards (CPH) and optimal survival tree (OST) models at 1, 2, 5 years and at the complete follow-up. The models are trained on 7 features (FSI) and 11 features (FSII).

|  | Model | Cohort | IBS FSI | IBS FSII |
| --- | --- | --- | --- | --- |
| IBS at 1 year | CPH | all | 0.0174 | 0.0174 |
| IBS at 2 years |  |  | 0.0353 | 0.034 |
| IBS at 5 years |  |  | 0.0677 | 0.0654 |
| overall IBS |  |  | 0.0962 | 0.0924 |
| IBS at 1 year | OST |  | 0.0169 | 0.0139 |
| IBS at 2 years |  |  | 0.035 | 0.0297 |
| IBS at 5 years |  |  | 0.0666 | 0.0573 |
| overall IBS |  |  | 0.0908 | 0.0797 |
| IBS at 1 year | CPH | MMRd | 0.0108 | 0.0092 |
| IBS at 2 years |  |  | 0.0318 | 0.0276 |
| IBS at 5 years |  |  | 0.0736 | 0.0665 |
| overall IBS |  |  | 0.1017 | 0.0957 |
| IBS at 1 year | OST |  | 0.0116 | 0.0081 |
| IBS at 2 years |  |  | 0.0339 | 0.0179 |
| IBS at 5 years |  |  | 0.0752 | 0.0416 |
| overall IBS |  |  | 0.1057 | 0.0728 |
| IBS at 1 year | CPH | NSMP | 0.009 | 0.0086 |
| IBS at 2 years |  |  | 0.017 | 0.0156 |
| IBS at 5 years |  |  | 0.0309 | 0.0306 |
| overall IBS |  |  | 0.0526 | 0.0505 |
| IBS at 1 year | OST |  | 0.0083 | 0.0084 |
| IBS at 2 years |  |  | 0.0175 | 0.0165 |
| IBS at 5 years |  |  | 0.0314 | 0.0301 |
| overall IBS |  |  | 0.0479 | 0.0471 |
| IBS at 1 year | CPH | p53ab | 0.0614 | 0.0593 |
| IBS at 2 years |  |  | 0.1014 | 0.0964 |
| IBS at 5 years |  |  | 0.1467 | 0.1466 |
| overall IBS |  |  | 0.1621 | 0.1561 |
| IBS at 1 year | OST |  | 0.0656 | 0.0559 |
| IBS at 2 years |  |  | 0.1053 | 0.0859 |
| IBS at 5 years |  |  | 0.1508 | 0.1185 |
| overall IBS |  |  | 0.1658 | 0.1326 |

### Additional ML models

We trained 9 survival analysis models on a full patient cohort. Supplementary Figure 1 displays model performance in the prediction of disease-specific survival in the full EC cohort using C-index as a metric. The cohort is preprocessed as indicated in the Materials and Methods section with three modifications. Firstly, we extended the FSII feature set with “leukocytes”, “hemoglobin”, “thrombocytes” variables, as well as thee CA125 and PR status indicators for a total of 16 variables. Secondly, we normalized the numerical variables to their z-scores, instead of winsorizing them. The z-score is calculated as follows:

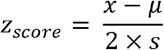

- *x* - numerical variable;
- *μ* - sample mean of *x*;
- *s* - sample standard deviation of *x*.

factor 2 is used in the denominator to make sure that the feature values are on the same scale as binary variables (0, 1), such that the distribution of transformed numerical variables has a mean of 0 and variance of 0.25 [71]. Thirdly, we use one-hot encoding to transform the categorical variables.

In addition to the CPH and OST models, seven tested models are:

- CoxNet model that extends linear Cox model by introducing L_1_ and L_2_ penalties to adjust for the impact of outliers [72];
- Gradient Boosting Machine (GBM) model is an ensemble method that fits shallow decision trees in a stage-wise manner, such that each subsequent tree is fitted on the residuals (errors) from the previous tree [73];
- Three models are based on Support Vector Machines (SVM) that perform survival modeling in the expanded feature space. We use SVMs with a linear kernel (linSVM), radial basis function kernel (rbfSVM) and minimal Lipschitz smoothness strategy linear kernel (lipSVM) [74];
- Random Survival Forest (RF) as implemented by Pölsterl [75];
- Stacked model that combines the above mentioned models by averaging their predictions.

We optimize hyperparameters of all ML models using 5-fold cross-validation, and then refit the models on the complete cohorts. The CI intervals are given as +/−1 standard deviation of the C-index calculated via 100 iterations of ordinary bootstrap with replacement.

**Supplementary Figure 1.**
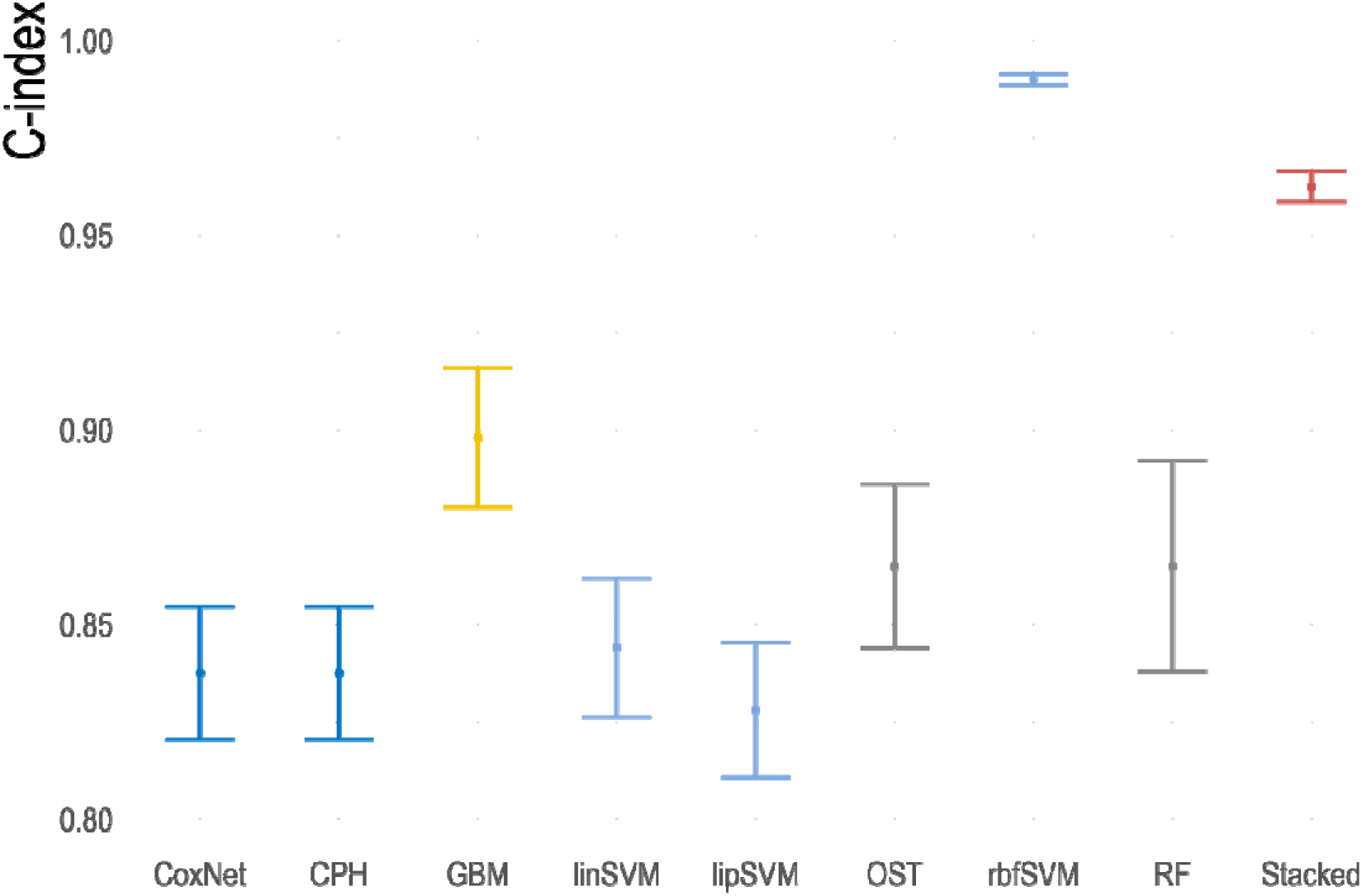
C-index of nine prognostic models trained on the full cohort (n = 700). The CI intervals are given as +/−1 standard deviation calculated via 100 iterations of ordinary bootstrap with replacement.

As seen in Supplementary Figure 1, rbfSVM leads to almost perfect C-index scores. Further, the GBM and Stacked models significantly outperform the CPH and OST models, which are the two main methods used in our work. Despite such impressive performance scores of the SVM, GBM and Stacked models, these methods are not easily interpretable, as such, they do not circumvent the main (from our point of view) limitation of ML methods in prognostic survival modeling - their “black box” nature. Difficulties in the model in interpretability are further compounded by insufficient performance reporting in the literature. These factors severely limit a more widespread use of ML and DL models in the clinical practice [18].

